# A testosterone metabolite 19-hydroxyandrostenedione induces neuroendocrine trans-differentiation of prostate cancer cells via an ectopic olfactory receptor

**DOI:** 10.1101/243204

**Authors:** Tatjana Abaffy, James R. Bain, Michael J. Muehlbauer, Ivan Spasojevic, Shweta Lodha, Elisa Bruguera, Sara K. O’Neal, So Young Kim, Hiroaki Matsunami

## Abstract

Olfactory receptor OR51E2, also known as a Prostate Specific G-Protein Receptor, is highly expressed in prostate cancer but its function is not well understood. Through *in silico* and *in vitro* analyses, we identified 24 agonists and 1 antagonist for this receptor. We detected that agonist 19-hydroxyandrostenedione, a product of the aromatase reaction, is endogenously produced upon receptor activation. We characterized the effects of receptor activation on metabolism using a prostate cancer cell line and demonstrated decreased intracellular anabolic signals and cell viability, induction of cell cycle arrest, and increased expression of neuronal markers. Furthermore, upregulation of neuron-specific enolase by agonist treatment was abolished in OR51E2-KO cells. The results of our study suggest that OR51E2 activation results in neuroendocrine trans-differentiation. These findings reveal a new role for OR51E2 and establish this G-protein coupled receptor as a novel therapeutic target in the treatment of prostate cancer.

**Significance:** Prostate cancer is the second most common cancer in men. Most deaths from prostate cancer are due to the progression of localized disease into metastatic, castration-resistant prostate cancer characterized by increased number of neuroendocrine-like cells. These neuroendocrine-like cells are non-proliferating, terminally differentiated cells. Olfactory receptor OR51E2, also known as a Prostate Specific G-Protein Receptor, is highly expressed in prostate cancer, and its expression correlates with disease progression. Here, we identify and validate novel endogenous ligands for this receptor. We show that activation of OR51E2 by newly-discovered prostate cancer-relevant agonists facilitates cellular transformation, resulting in neuroendocrine trans-differentiation, a characteristic phenotype of castrate resistant prostate cancer. Our results establish this G-protein coupled receptor as a novel and therapeutic target for castration-resistant prostate cancer.

**Highlights:**

- Discovery of novel agonists for olfactory receptor OR51E2/PSGR highly relevant to prostate cancer pathology
- Activation of OR51E2 receptor by agonist *N*-acetyl-*N*-formyl-5-methoxykynurenamine (AFMK) results in release of 19-hydroxyandrostenedione (19-OH AD) from the prostate cancer cells indicating its endogenous production
- Activation of OR51E2 receptor by 19-OH AD, AFMK, and propionic acid decreases anabolic and proliferative signals
- Activation of OR51E2 receptor by 19-OH AD and AFMK increases markers specific for neuroendocrine trans-differentiation (NEtD)
- Ablation of the OR51E2 gene in prostate cancer cells treated with agonist 19-OH AD significantly reduces neuron-specific enolase

## INTRODUCTION

G-protein coupled receptors (GPCRs) have emerged as important factors in tumor growth and metastasis ^1–3^. Several GPCRs, such as the 5HT1c serotonin receptor ^4^, the M1, M3, and M5 muscarinic receptors ^5^, and the α1B-ADR adrenergic receptor ^6^, can function as oncogenes when persistently activated. These GPCRs, which are normally expressed in fully differentiated, post-mitotic neuronal cells, are able to induce cellular oncogenic transformation when introduced to an ectopic environment of proliferating cells and activated by agonist ^7,8^. In addition to oncogenes and tumor-suppressor genes essential for cancer initiation and progression, autocrine and/or paracrine secretion of GPCR-activating molecules and their downstream signaling events affect tumor growth, survival, and metastasis ^1,9–14^.

Olfactory receptors (ORs) are the largest family of GPCRs present in the olfactory epithelium but are also found in various ectopic or non-olfactory locations such as prostate, heart, placenta, embryo, erythroid cells, spleen, kidney, gut, tongue, and carotid body ^15–17^. Some ectopic ORs also play roles in chemotaxis^18^, muscle regeneration^19^, blood pressure regulation^20^, and hypoxia response^21^. OR51E2, or Prostate Specific G-protein Receptor (PSGR), is one of the most highly conserved and broadly expressed ectopic ORs ^22–24^. OR51E2 is present in healthy prostate tissue and shows significantly increased expression in prostate intraepithelial neoplasia (PIN), prostate adenocarcinoma (PC), and castration-resistant prostate cancer (CRPC) ^25–32^.

### What is the role of OR51E2 in prostate cancer?

The majority of prostate tumors start as androgen-dependent adenocarcinomas. As localized cancer progresses to a metastatic state, the number of neuroendocrine (NE)-like cells increases, contributing to the development of a highly aggressive form of castrate-resistant prostate cancer (CRPC) known as neuroendocrine prostate cancer (NEPC)^33,34^. Many clinical studies have demonstrated a correlation between neuroendocrine trans-differentiation (NEtD) and PC progression with poor prognosis ^35^. Tumor-derived NE-like cells are localized in tumor foci and are non-proliferating, terminally differentiated cells rich in serotonin and positive for NE markers, including neuron-specific enolase (NSE) and chromogranin A (CGA) ^35^.

The molecular mechanism underlying development of a neuroendocrine phenotype in PC is not fully understood. In the androgen-dependent PC cell line LNCaP, serum deprivation and agents that increase cAMP also increase expression of NEtD markers and genes indicative of neuronal phenotype ^36^.

In the olfactory system, ORs signal via the canonical cAMP pathway^37^, and several reports have indicated cAMP-mediated signaling for ectopic ORs^38–40^. Furthermore, high expression of the OR downstream targets adenylate cyclase 3 and Gaolf was recently identified in prostate tissue, supporting the role of a cAMP-mediated pathway in ectopic OR activation^22,41^. We hypothesized that agonist-mediated activation of OR51E2 increases cAMP and facilitates cellular transformation, resulting in NEtD. Thus, ectopic expression of this GPCR in proliferating cells and ligand-dependent activation could enable this receptor to function as an oncogene.

Recently, it has been demonstrated that overexpression of OR51E2/PSGR in a PSGR-Pten (Δ/Δ) mouse model accelerates PC development and progression ^42^. Furthermore, β-ionone, an agonist for OR51E2, decreased proliferation and increased invasiveness of human PC cells ^43–45^.

In this paper, we aimed to identify biologically relevant OR51E2 ligands using a combination of *in silico* investigations and experimental validation, and we also set out to study the effects of these ligands on androgen-dependent LNCaP cells. Currently identified OR51E2 agonists include the short-chain fatty acids acetate and propionate ^20,46^, steroid derivatives, β-ionone ^43^, and lactate ^21^. However, it is not known whether activation of OE51E2 by endogenous ligands is involved in PC pathogenesis.

Here, we virtually screened >2,500 metabolites, experimentally validated 55 of these candidates *in vitro,* and ultimately identified 24 new agonists and 1 antagonist for the human OR51E2 receptor. Among the agonists, we identified 19-hydroxyandrostenedione (19-OH AD)—which is synthesized by aromatase, an enzyme highly expressed in NEPC and CRPC—and *N*-acetyl-*N*-formyl-5-methoxykynurenamine (AFMK), a tryptophan and melatonin metabolite. We detected endogenous production of 19-OH AD in the LNCaP cells stimulated with AFMK agonist. The identity of the newly discovered agonists, as well as significant differences in metabolomics signatures in agonist-stimulated PC cells indicative of non-proliferation, prompted us to further investigate their effects on cell viability, cell cycle status, and several NE markers to examine if OR51E2 receptor activation drives NEtD.

## RESULTS

### *In Vitro* Validation of Ligands Predicted *in Silico*

A structural model of OR51E2 was made *in silico* using Modeller (see Materials and Methods for details). To experimentally validate our virtual library screening (VLS) predictions, we used an *in vitro* heterologous expression system ^47^ in which Hana 3A cells transfected with the OR51E2 receptor were stimulated with the top candidate ligands. Responses were subsequently measured with a cAMP-mediated luciferase reporter gene assay. First, a small library of 33 compounds identified as PC-associated metabolites was selected from the Human Metabolome Database HMDB ^48^. These compounds were identified from previous reports ^49–51^. Additional compounds previously identified as OR51E2 agonists were also included: 1,4,6-androstatriene-17-β-ol-3-one, 1,4,6-androstatriene-3,17-dione, 6-dehydrotestosterone, and β-ionone^43^. Thus, a total of 37 compounds were used for the initial, small-scale VLS (**Table S4**). Two scoring functions, Score and mfScore, were used to predict the best binders. The top 9 compounds from each score list (italicized in **Table S4**) were then tested *in vitro,* and the following metabolites were identified as novel agonists for OR51E2 (bold and italics): bradykinin, kojibiose, glycylglycine, L-histidinol N-acetylglutamic acid, and D-alanyl-D-alanine. We also confirmed the previously reported agonist 1,4,6-androstatriene-3,17-dione (**Figure S2**)^43^.

Next, a larger library of 2,511 human metabolites from the HMDB was selected for virtual screening and docking into the receptor pocket (**Figures 1A and 1B**). Here, the top potential ligands (in italics, **Tables S1 and S2**) were tested *in vitro* using a biologically relevant concentration range reported in the HMDB, and concentration-response curves were subsequently produced. In total, 55 compounds were tested (9 and 46, from the smaller and larger screens, respectively), and 24 agonists (**Figures 1C, and S2 and S3**) and 1 antagonist for OR51E2 (**Figure 1C**) were identified. In each experiment, OR51E2-expressing Hana3A cells were also stimulated with a known agonist, 1 mM propionic acid (PA), so we were able to compare the efficacy of each metabolite relative to PA. Furthermore, potency values (EC_50_) were determined (**Table 1**). Glycylglycine was the most efficient agonist, while L-histidinol was the most potent. Diverse metabolites were discovered as novel agonists for OR51E2. Concentration-response curves for metabolites from the large VLS screen that did not activate the receptor are presented in **Figure S4**.

**Table 1.** Potency and efficacy of 24 newly discovered OR51E2 agonists with their associated HMDB and CAS identifiers. 1,4,6-androstatriene-3,17-dione (**Figure S2**) was not included, as it is one of the previously identified compounds. Maximal concentration of each chemical used to measure efficacy in comparison to the response of 1 mM propionic acid (PA) was also indicated. *Imidazolone full name = 4-methyl-2,3-dihydro-1H-imidazol-2-one.

| NAME | HMDB | CAS | EC<sub>50</sub> (potency)<br>[M] | Max conc.<br>used | Efficacy<br>(relative to<br>1 mM PA) |
| --- | --- | --- | --- | --- | --- |
| D-Alanyl-d-alanine | HMDB03459 | 923-16-0 | 1.40E-05 | 3.16 mM | 1.505 |
| AFMK | HMDB04259 | 52450-38-1 | 1.20E-05 | 3.16 mM | 1.483 |
| Gamma-CEHC | HMDB01931 | 178167-77-6 | 6.40E-09 | 10 µM | 1.286 |
| Hydroxypyruvic acid | HMDB01352 | 1113-60-6 | 4.20E-07 | 316 µM | 1.100 |
| Adenosine 2',3'-c-phosphate | HMDB11616 | 634-01-5 | 2.60E-08 | 3.16 µM | 1.025 |
| Palmitic acid | HMDB00220 | 57-10-3 | 9.80E-09 | 1 mM | 0.927 |
| L-Glyceric acid | HMDB06372 | 28305-26-2 | 1.90E-09 | 1 mM | 0.898 |
| 19-OH AD | HMDB03955 | 510-64-5 | 1.50E-10 | 10 µM | 0.890 |
| Androstanedione | HMDB00899 | 846-46-8 | 7.90E-10 | 100 µM | 0.888 |
| N-Acetylglutamic acid | HMDB01138 | 1188-37-0 | 2.30E-10 | 10 µM | 0.879 |
| Kojibiose | HMDB11742 | NA | 1.00E-06 | 316 µM | 0.790 |
| Bradykinin | HMDB04246 | 58-82-2 | 1.30E-09 | 100 µM | 0.762 |
| Imidazolone* | HMDB04363 | 1192-34-3 | 7.60E-12 | 10 µM | 0.678 |
| Pelargonidin | HMDB03263 | 134-04-3 | 4.20E-10 | 100 µM | 0.621 |
| Glycine | HMDB00123 | 56-40-6 | 5.80E-08 | 1 mM | 0.613 |
| 2-Ketoglutaric acid | HMDB00208 | 328-50-7 | 5.50E-09 | 1 mM | 0.594 |
| Urea | HMDB00294 | 57-13-6 | 2.30E-08 | 10 mM | 0.580 |
| L-Histidinol | HMDB03431 | 4836-52-6 | 3.50E-11 | 100 µM | 0.578 |
| 8-Hydroxyguanine | HMDB02032 | 5614-64-2 | 4.40E-13 | 100 nM | 0.570 |
| 2-Pyrrolidinone | HMDB02039 | 616-45-5 | 1.90E-09 | 100 µM | 0.525 |
| Epitestosterone | HMDB00628 | 481-30-1 | 6.90E-10 | 10 µM | 0.477 |
| Estriol | HMDB00153 | 50-27-1 | 5.30E-05 | 10 µM | 0.344 |
| Tetrahydrocurcumin | HMDB05789 | 36062-04-1 | 5.70E-07 | 316 µM | 0.284 |

**Figure 1.**
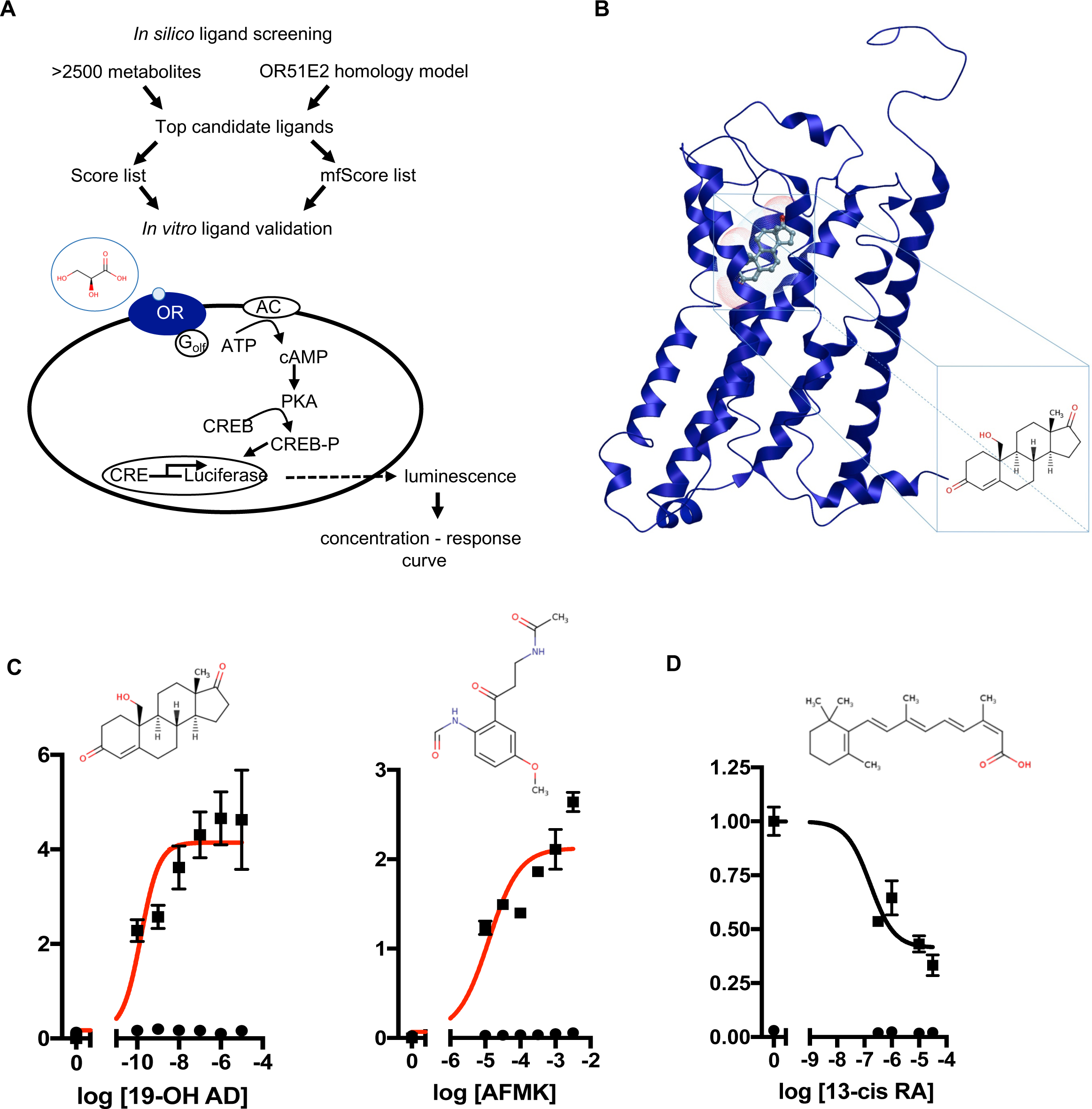
Discovery of novel endogenous metabolite-ligands for OR51E2. **A.** Study design. **B.** Homology model of OR51E2 with 19-hydroxyandrost-4-ene-3,17-dione (19-OH AD) docked into the receptor pocket. **C.** Concentration - response curves for 19-OH AD and AFMK (red) and their structures. **D.** Concentration - response curve for antagonist 13-cis retinoic acid (13-cis RA). N=3, mean ± SEM.

Some of the newly discovered agonists are: 19-hydroxyandrostenedione (19-OH AD), a hypertensive steroid (vasopressor) secreted by the adrenal gland ^52–54^, an intermediate in estrogen synthesis from testosterone ^55^ also found in porcine testes ^56^ and rat ovarian granulosa cells ^57^; acetyl-N-formyl-5-methoxykynurenamine (AFMK), a melatonin and kynurenamine metabolite previously reported to be abundant in aggressive PC ^58^; estradiol ^59^; adenosine-2,3-cyclic phosphate, a positional isomer of the second messenger 3’,5’-cAMP ^60^; 8-hydroxyguanine, a marker of DNA damage; α-ketoglutaric acid, an intermediate in the citric acid cycle; urea; glycine; and palmitic acid.

Because *in situ* estrogen production is an important factor in prostate carcinogenesis, and since the expression of aromatase, the enzyme that synthetizes estrogens from androgens, is increased 30-fold in PC, we decided to further examine the presence and production of 19-OH AD in LNCaP cells^61,62^.

### Endogenous 19-OH AD production upon OR51E2 activation with AFMK

We have developed a liquid chromatography/mass spectrometry (LC-MS/MS) assay for the measurements of 19-OH AD in the cell media. LNCaP cells were stimulated with newly discovered agonist, 250 μM AFMK, for 3 days, and the 19-OH AD was measured in the cell medium (**Figure 2**). We used CD-phenol free RPMI1640 medium (**Figure 2A**) and regular RPMI1640 medium (**Figure 2B**) to estimate production of 19-OH AD. Three times more 19-OH AD was detected when cells were stimulated in CD-phenol free medium (0.83 *vs.* 0.27ng/mL). No 19-OH AD was detected in unstimulated cells (dotted lines in **Figures 2A and 2B**).

**Figure 2.**
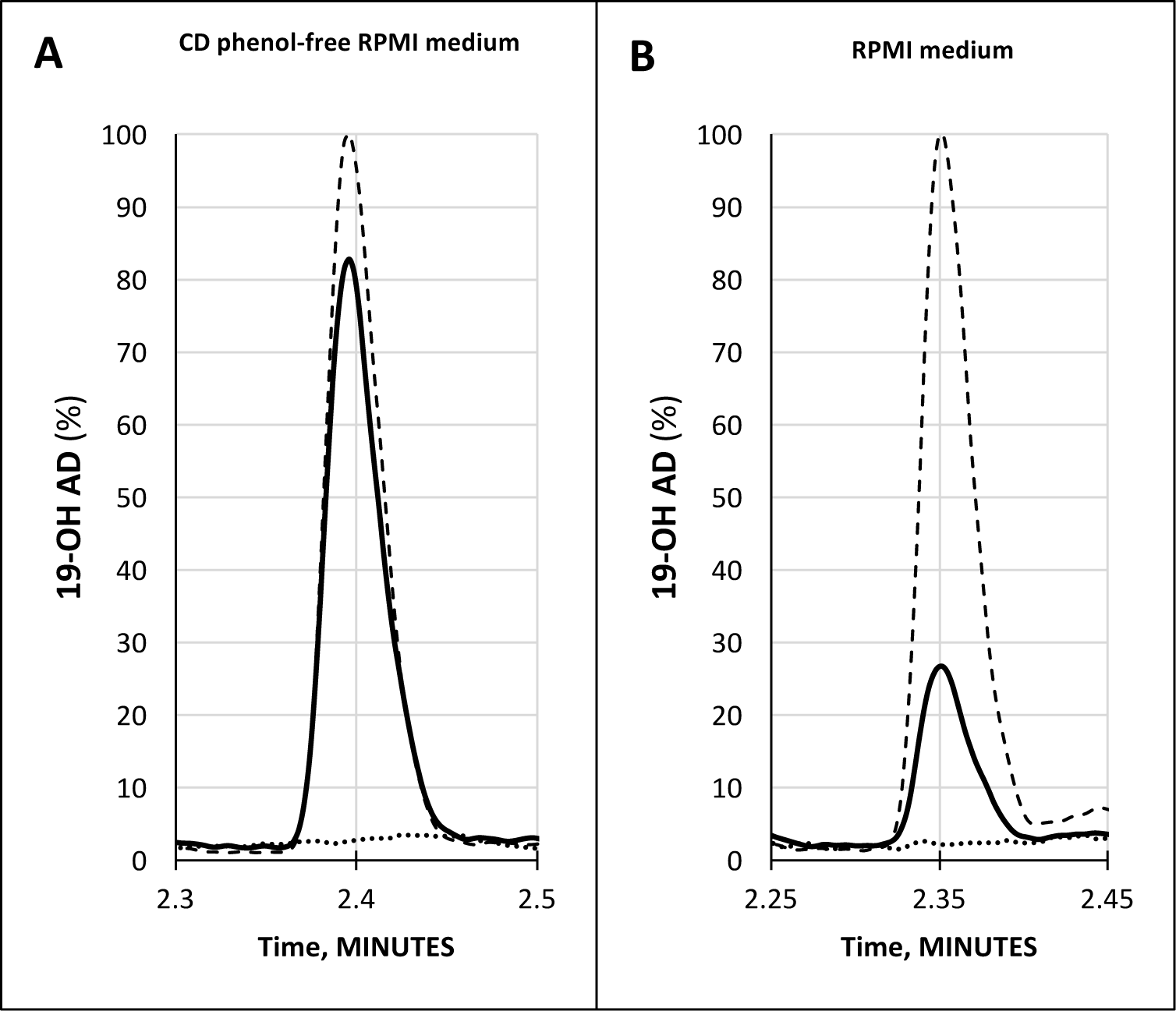
Quantification of 19-OH AD by LC-MS/MS in the medium from AFMK-stimulated LNCaP cells. **(A)** CD phenol-free RPMI medium with 250 μM AFMK (solid line), 1 ng/mL of 19-OH AD-standard spike (dashed line); estimated concentration of 19-OH AD is 0.83 ng/mL medium. **(B)** RPMI medium + 250 μM AFMK (solid line), 1 ng/mL 19-OH AD of standard spike (dashed line); estimated concentration of 19-OH AD is 0.27 ng/mL. Dotted lines at the bottom in A and B are media only, with no AFMK. Curves are normalized to 1ng/mL 19-OH AD.

### Metabolomic Signatures of LNCaP Cells Treated with Selected OR51E2 Agonists

In addition to the newly discovered agonists 19-OH AD and AFMK, we also selected the previously identified OR51E2 agonist propionic acid (PA) for metabolomics analysis ^46^. Cells were incubated with 100 nM 19-OH AD, 250 μM AFMK, and 1 mM PA for 72 h, and non-targeted metabolomics analysis was performed to identify metabolites that differed significantly in the cell lysate and conditioned medium (CM). Agonist treatment resulted in pronounced intra- and extra-cellular changes in metabolomic signatures, as seen in heat maps (**Figures S5 and S6**, respectively). Differentially expressed features/metabolites identified in each group using the t-test (*P* < 0.05) and fold change (2-fold or greater) are presented in **Figures S5-S10** and **Tables S5-S10**.

The top 15 differentially expressed extracellular metabolites are presented in **Figures 3A, 3B**, and 3C. All 3 agonists also produced robust intracellular decreases in amino acids, especially serine and threonine, and also 2 glycolytic intermediates: glucose-6-P and fructose-6-P (**Figure 3D, 3E and 3F**). Furthermore, agonist treatment resulted in decreases in both methionine and glycine levels. In addition to decreased lactic acid, significant decreases were noted in fumaric, malic, and succinic acids, all intermediates of the TCA. Interestingly, we detected an increased level of phosphoenolpyruvate (PEP) in all agonist-treated samples (fold change analysis, **Figures S7B, S8B and S9B**). We also observed decreased levels of myoinositol, inosine, adenosine, asparagine, aspartate, and guanosine, which have been previously reported as being depleted in metastatic PC tissue ^49^, indicating that activation of OR51E2 by agonists in LNCaP cells produces metabolic signatures similar to those observed in human metastatic tissues. Agonist treatment also reduced levels of intracellular urea and spermine/spermidine, in accordance with recently reported data showing reduced spermine during malignant transformation ^63^. Accordingly, we also detected decreased intracellular levels of ornithine, a precursor of polyamines.

**Figure 3.**
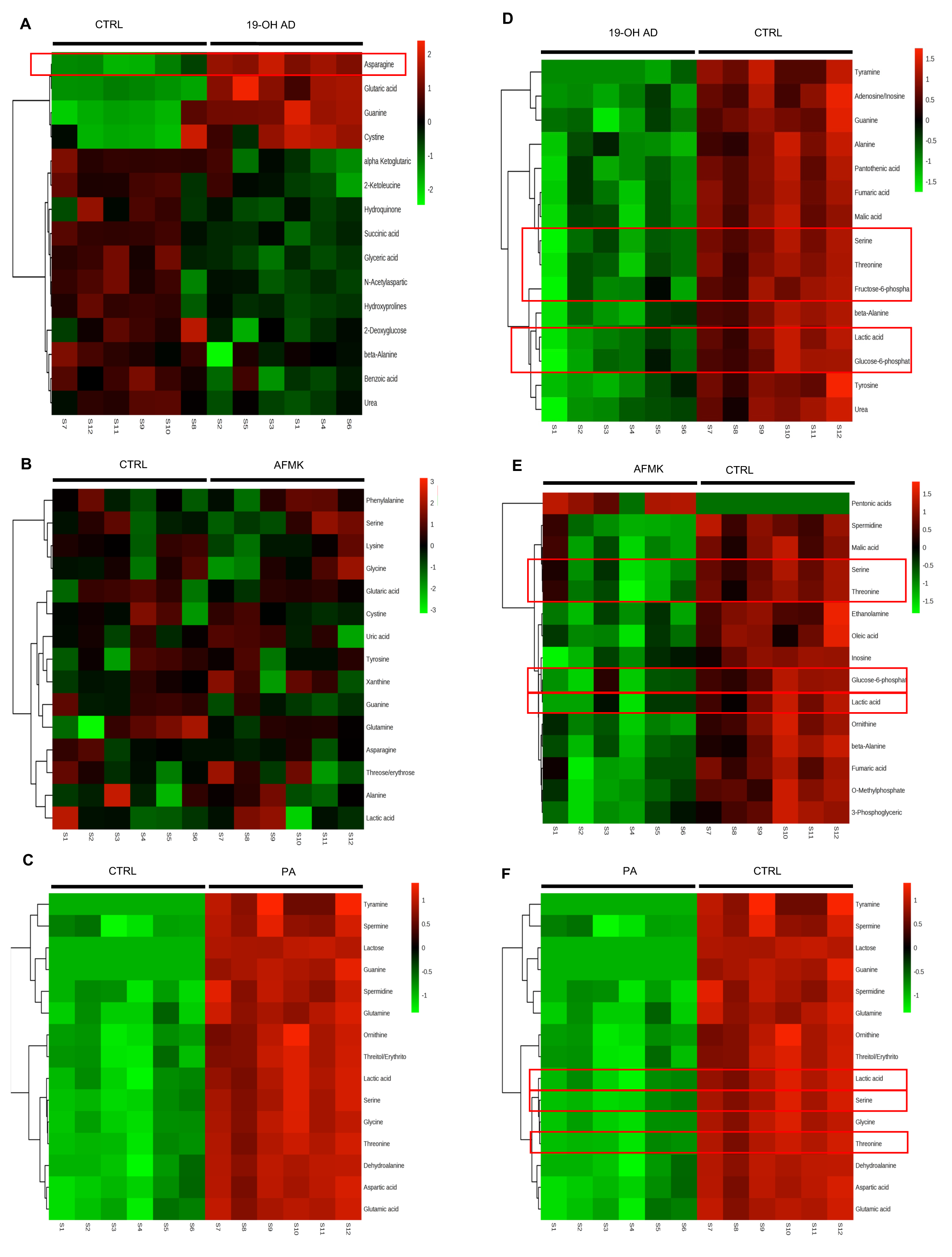
Agonist treatment of LNCaP cells results in robust metabolomic signatures. The top 15 extracellular metabolites identified after stimulation with **A.** 19-OH AD, **B.** AFMK, and **C.** PA. The top 15 intracellular metabolites identified after stimulation with **D.** 19-OH AD, **E.** AFMK, and **F.** PA. Heatmaps are based on the Pearson correlation analysis (Ward) and indicate annotated metabolites identified by t-test (*P <* 0.05, FDR <0.1, n = 6). Columns correspond to the samples treated with agonists (S1–6) and control (S7–12), and rows correspond to annotated metabolites.

Fold change analysis of the CM revealed significantly increased glutamine (25-, 11-, and 51-fold, for 19-OH AD, AFMK, and PA, respectively), indicating that agonist-treated cells do not have an increased demand for glutamine as highly proliferating cancer cells usually do (**Figures S10B, S11B, and S12B**). Thus, these results argue for a non- or a low-proliferative phenotype. Decreased levels of docosanoic and decanoic acid and increased levels of asparagine were also prominent in the CM in all 3 treatments (**Figure 4G**). The pathways most affected, as identified by MetaboAnalyst, were the serine, threonine, and glycine; alanine, aspartate, and glutamine; ketogenesis; arginine and proline; and beta-alanine metabolic pathways (**Figure 4F**).

**Figure 4.**
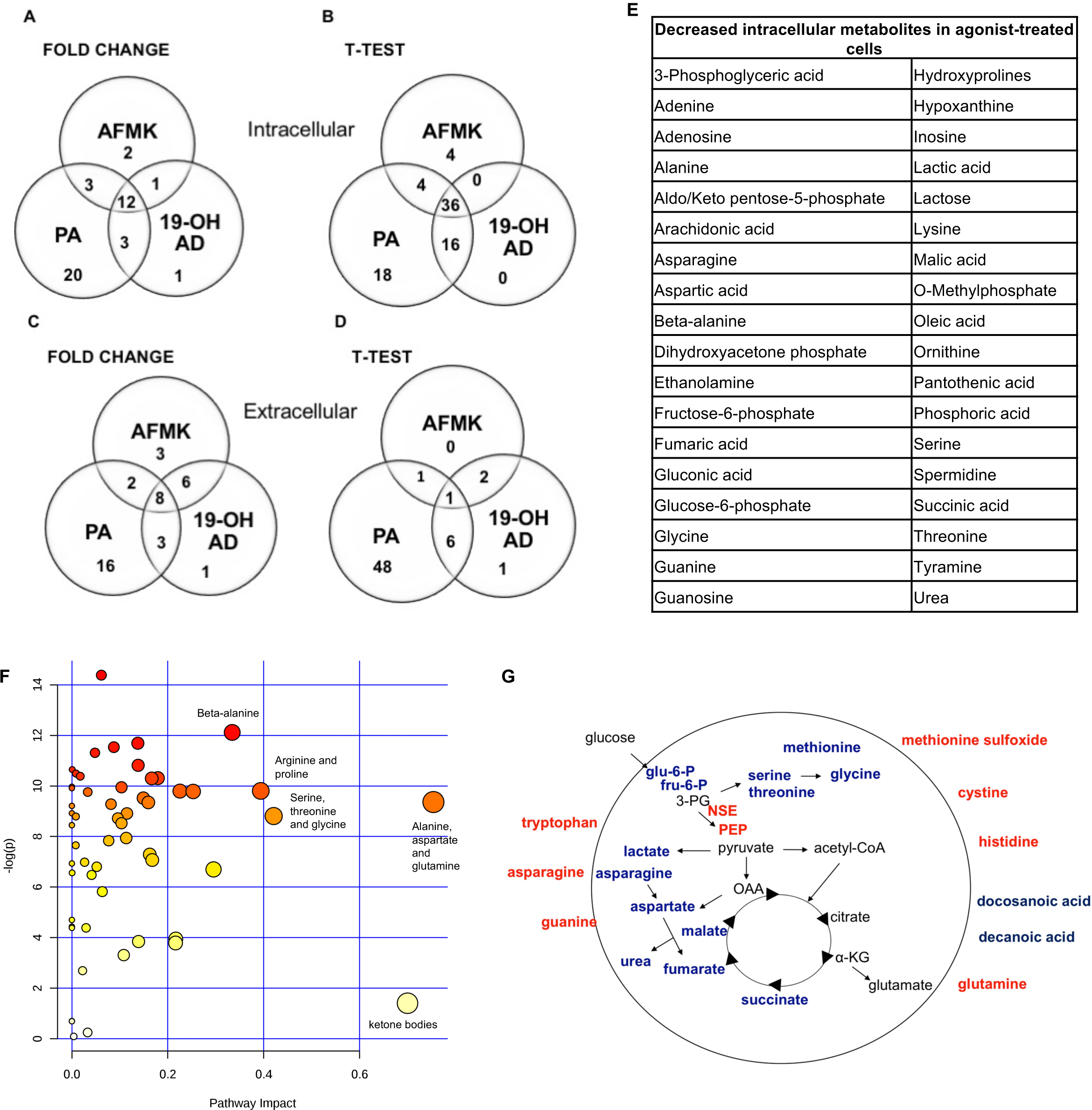
Most pronounced metabolic changes induced by activation of OR51E2 with selected agonists. Venn diagrams show the set of annotated intracellular metabolites differentially expressed by **(A)** fold change and **(B)** t-test analysis, and the set of extracellular metabolites identified by **(C)** fold change and **(D)** t-test. **E.** In total, 36 annotated intracellular metabolites significantly decreased in all 3 treatments (see **3B). F.** Pathway analysis. The most pronounced pathways in 19-OH AD-treated cells. Pathways are displayed as circles, and the color and size of each circle is based on its *P* value and pathway impact value, respectively. The top-right area indicate the most significant changes in metabolites. **G.** Schematic representation of cellular metabolites. Significantly increased metabolites are in bold red, and significantly decreased are in bold blue (PEP - phosphoenol pyruvate, OAA - oxalacetate, 3PG – 3-phosphoglycerate, α-KG - α-ketoglutarate). Increased NSE (neuron-specific enolase) catalyzes the formation of PEP and is also indicated in red.

### Time-dependent Modulation of Cellular Proliferation

Our metabolomics results indicated reduced capacity for anabolic reactions in LNCaP cells following receptor activation, which prompted us to further analyze the effects of OR51E2 agonists on cellular proliferation. Moreover, because melatonin reduces cell proliferation, we examined whether AFMK, a melatonin metabolite, also reduces or inhibits cell proliferation ^64^. LNCaP cells were stimulated with various concentrations of 19-OH AD and AFMK for 4 days and analyzed every 24 hours using an ATP viability assay. At day 4, both agonists significantly decreased the number of viable cells when compared to the control, non-stimulated cells (**Figure 5A**). This effect was dependent on cell seeding density; at higher plating densities (40% to 50%), the effect was not observed.

**Figure 5.**
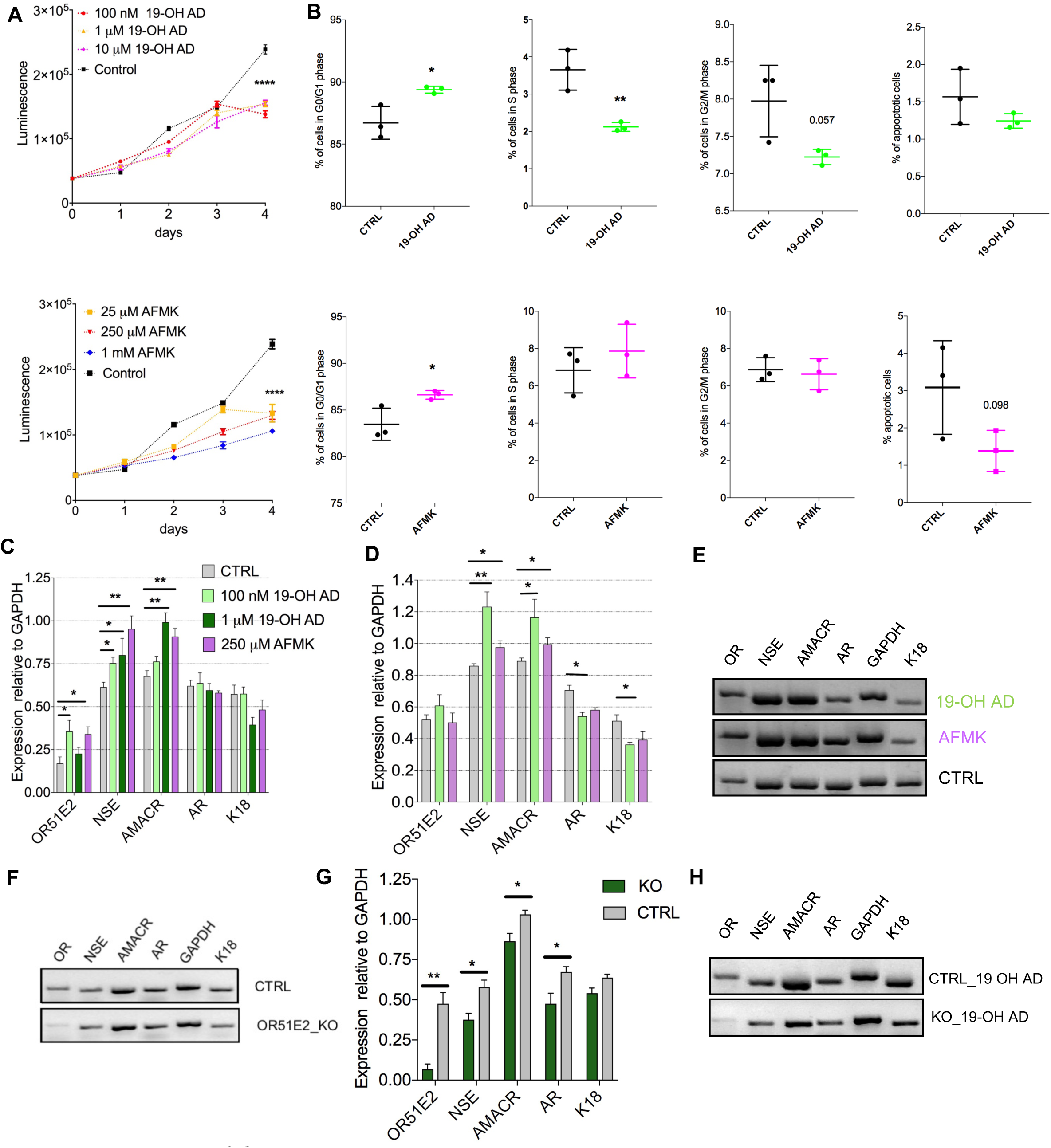
Activation of OR51E2 receptor by selected agonists induces a neuroendocrine phenotype. **A.** Cell viability assay at various indicated concentrations of 19-OH AD and AFMK. Cell viability correlates with luminescence signal. Statistical significance at day 4 with 100 nM 19-OH AD and 250 μM AFMK is \*\*\*\**P* <0.0001. **B.** Cell cycle analysis after incubation with 100 nM 19-OH AD and 250 μM AFMK for 7 and 3 days, respectively. **C.** Transcript levels of markers after agonist stimulation for 3 days; N = 3 to 6, unpaired t-test, **P <0.01. *P <0.05 **D.** Transcript levels of makers after stimulation with agonists for 12 days, N= 3 to 6, unpaired t-test, \*\**P* <0.01. \**P* <0.05 **E.** Representative gel image of data presented in D. **F.** Markers in the OR51E2-knockout (KO) and control cells (CTRL). **G.** Transcript levels of markers relative to GAPDH after 3 days stimulation with 1 μM 19-OH AD in OR51E2-KO LNCaP cells. N=4, mean ± SEM. \*\**P* <0.01. \**P* <0.05.

### Cell Cycle Arrest

Next, we determined if the decrease in cell viability during agonist stimulation is attributable to increased apoptosis or cell-cycle arrest. Cells were treated with various concentrations of agonists for 3 and 7 days. Both agonists increased the fraction of cells in the G0/G1 phase and decreased the number of apoptotic cells (**Figure 5B**). Our results are in agreement with previous reports that the majority of cells with a neuroendocrine-like phenotype show signs of resistance to apoptosis ^65^.

### Neuroendocrine Markers

To assess the effect of selected OR51E2 agonists on NEtD, we analyzed transcript levels after 3 and 12 days of stimulation of the following neuroendocrine, epithelial, and receptor genes: NSE; α-methylacyl-CoA racemase (AMACR); keratins 5, 8, and 18; voltage-gated Ca channel a1H (Cav3.2); androgen receptor (AR); and OR51E2 receptor. NSE was used to specifically identify NEtD status, and although LNCaP (being a cell line established from lymph-node metastatasis) cells already express low levels of NSE, treatment with OR51E2 agonists significantly increase levels of the NSE transcript (**Figures 5C, 5D, and 5E**). These results correlate well with the increased intracellular level of PEP detected in the metabolomic analysis (**Figures S7B, S8B and S9B**), since the glycolytic enzyme enolase catalyzes PEP synthesis. α-AMACR is an enzyme essential for isomerization of branched-chained fatty acids and is present at very low levels in healthy prostate and increased in PC and NE-like cells ^66^. Since NE-like tumor cells express AMACR, we investigated whether activation of OR51E2 also increased AMACR levels. Indeed, AMACR levels were significantly increased after 3 and 12 days of agonist stimulation (**Figures 5C, 5D, and 5E**).

Furthermore, 19-OH AD decreased the AR transcript after 12 days, and although AFMK also showed a similar trend, it did not reach statistical significance (**Figure 5D**). Normal basal prostate epithelial cells are positive for K5, and expression of K5 is also associated with the epithelial-to-mesenchymal transition during tumor progression and metastasis ^67^. K5 and K8 were not detected in the agonist-stimulated cells (data not shown). A significant decrease in K18, a luminal secretory marker, was found after 12 days of stimulation with 100 nM 19-OH AD (**Figure 5D**). Although Ca+^2^ entry through the voltage-gated calcium channel α1H (Cav3.2) was previously reported to be involved in NEtD of LNCaP cells when cultured in steroid-free conditions, we did not detect changes in its transcript levels (data not shown) ^68^.

### OR51E2 Knock-out Confirms NE-like Phenotype Upon Receptor Activation

To confirm receptor involvement in the agonist-mediated increase of NSE, OR51E2 was deleted using a CRISPR-Cas9 method (**Figure 5F**). We designed 3 gRNAs to target Cas9 to the OR51E2-gene and generated a lentiviral sgRNA-Cas9 vector to deliver gRNA into cells. The efficiency of each gRNA was measured, and we observed that by using sgRNA #1, OR51E2 was abrogated in 80% of cells. These OR51E2-knockout cells were exposed to 1 μM 19-OH AD for 3 days and analyzed for the presence of specific markers (**Figures 5G and 5H**). A statistically significant decrease of NSE in OR51E2-knockout cells in comparison with control cells was observed (from 0.579 ± 0.043 in control to 0.377 ± 0.04, *P* < 0.05, n = 4 biological replicates, **Figure 5G**), confirming that increased NSE during stimulation with agonists is at least partially a receptor-mediated phenomenon (**Figures 5C and 5D**).

## DISCUSSION

As the number of ectopic olfactory receptors associated with diverse pathological states continues to increase ^15,17^, the implications and significance of these receptors will be greatly enhanced by receptor “deorphanization” (i.e., defining the ligands). Previously, we successfully identified novel ligands for mouse OR using a similar *in silico* approach with VLS ^69^. Here, we present a highly successful approach of combining *in silico* and *in vitro* analyses to identify novel biologically relevant ligands for the human ectopic OR OR51E2. This method can be used to elucidate ligand specificities of other ectopic ORs. Once identified, these new ligands can help define the role and function of ORs in cancer and other diseases.

Among the newly discovered metabolites identified as OR51E2 agonists, several were previously reported to be associated with PC, including bradykinin, kojibiose, glycylglycine, *N*-acetylglutamic acid, and D-alanyl-D-alanine ^49^. Thus, our results indicate that these metabolites are likely endogenous agonists. New agonists with known biological roles were also discovered: epitestosterone, known to be a major metabolite of androstenedione and testosterone ^70^; and androstanedione (also known as 5α-androstane-3,17-dione), an intermediate in steroid synthesis^71^.

In addition to these agonists, we also identified a previously under-reported metabolite of the complex steroid biosynthetic network, 19-hydroxyandrost-4-ene-3,17-dione (19-OH AD) ^55,72,73^. 19-OH AD is produced by aromatase P450 (CYP19A1), which catalyzes the irreversible aromatization of the androgens androst-4-ene-3,17-dione and testosterone and their consequent conversion to estrogens (http://www.brenda-enzymes.org/enzyme.php?ecno=1.14.14.14). We detected this testosterone metabolite in agonist-stimulated prostate cancer cells. These results demonstrate that 19-OH AD is actively produced by cancer cells when the OR51E2 receptor is activated. Thus, we demonstrate that 19-OH AD is an endogenous agonist produced by activation of OR51E2 in prostate cancer cells.

Aromatase is increased 30-fold in metastatic PC ^62^, and aromatase-knockout mice have a reduced incidence of PC following exposure to testosterone and estrogen, indicating that aromatase metabolites, mainly 19-OH AD and estradiol, are likely involved in prostate carcinogenesis. Results from our study demonstrate that 19-OH AD is a potent OR51E2 agonist (EC_50_= 1.5^−10^ M) and support the notion that increased *in situ* estrogen production via 19-OH AD is an important factor in PC ^61^.

AFMK is a metabolite of melatonin ^74^. Previous studies demonstrated that melatonin reduces proliferation of LNCaP cells, leading to NEtD, and the phenotype was not reversed by melatonin receptor antagonists, suggesting that additional receptors may be mediating this process ^64,75^. Our results demonstrate that OR51E2 is a receptor for AFMK, a melatonin metabolite, and although its EC_50_ is in the μM range, much higher than reported blood concentrations (<65 pM) ^76^, the local tissue concentration may reach μM and mM levels as has been recently shown for keratinocytes ^74^. An additional source of AFMK might be a tryptophan metabolic pathway ^77^. Tumors produce high levels of tryptophan and kynurenic acid metabolites ^78^. Significant amplification of tryptophan-2,3-dioxygenase (EC 1.13.11.11) TDO2, which catalyzes the oxidation of L-tryptophan to *N*-formyl-L-kynurenamine, was observed in NEPC and PC ^79,80^. Thus, in more advanced stages of PC, AFMK production may be increased via this tryptophan metabolic pathway. Recently, this pathway was shown to regulate the immunosuppressive microenvironment of various tumors^81^.

We also identified bradykinin as an agonist for OR51E2. In general, kinins are released during the inflammatory reaction and they are involved in angiogenesis and tumorigenesis ^82,83^. Our results indicate that the activation of OR51E2 by bradykinin can also happen in the early stages of PC, when the inflammatory milieu is predominant. Prostatic secretions of PC patients have elevated levels of human kallikrein 2 ^84^, which produces bradykinin and thus stimulates proliferation of androgen-independent PC cells in later stages of PC ^85^.

The OR51E2 antagonist 13-cis RA is an endogenous component of human serum, and many of its actions can be explained by isomerization to all-trans RA and 9-cis RA, which both act via retinoid receptors. However, since 13-cis RA does not have potent gene regulatory activity, additional pathways via membrane receptors have been proposed to explain its pharmacological and anti-inflammatory actions ^86^. Our results demonstrate that 13-cis RA acts via the OR51E2 receptor when expressed heterologously.

OR51E2 receptor activation by 19-OH AD, AFMK, and PA induced pronounced metabolic reprograming of LNCaP cells, with the most significant changes being decreased intracellular serine and threonine levels. Because metabolism of these amino acids includes one-carbon metabolism, which provides cofactors for biosynthetic reactions in highly proliferating cells, intracellular depletion may indicate a general decrease in anabolic reactions ^87^. Furthermore, an intracellular decrease in aspartate, which is normally required for protein, purine, and pyrimidine synthesis, and an increase in the CM indicate that agonist-activated LNCaP cells are not preparing for proliferation. We also detected decreased intermediates of glycolysis (glucose-6-phosphate and fructose-6-phosphate) in activated cells. Agonist treatment decreased intracellular lactate, suggesting a slower rate of glycolysis. An intriguing result was the increased intracellular level of phosphoenolpyruvate (PEP). We also found increased NSE transcription for the glycolytic enzyme enolase, which catalyzes the formation of PEP, indicating predominance of the PEP-forming reaction.

NSE is not only a marker of neuronal differentiation and maturation characteristic of neurons and neuroendocrine cells ^88^, but it also has an important role in synaptogenesis ^89^ and is reported to be stable in the high-chloride environment characteristic of neurons, in which it reaches a concentration of 2% to 3% of the soluble protein ^90^. Taken together, these results indicate that receptor activation results in a neuronal-like phenotype of LNCaP cells.

Cystine was increased in the medium after 19-OH AD and AFMK treatment (Tables S12 and S13). In healthy cells, cystine is transported into cells and reduced to cysteine, which can then be utilized for synthesis of gluthatione, a protective antioxidant. As a consequence of rapid cell growth during tumorigenesis, the production of reactive oxygen species increases, providing a proliferative signal for glutamine to enter the cell and, after deamidation, condense with cysteine to form a precursor of glutathione. However, in our experiments, the medium, but not the cells, showed increased levels of glutamine and cysteine, indicating a reduction in protective oxidative and proliferative signals in agonist-stimulated cells. The alanine/aspartate/glutamine pathway is the most affected biochemical pathway during NEtD of LNCaP cells induced by steroid-reduced medium, which corroborates our results ^91^ (**Figure 4F**). In cancer-associated fibroblasts, asparagine and aspartate are involved in glutamine synthesis^92^, and our experiments showed decreased intracellular levels of these amino acids, suggesting increased use for intracellular synthesis of glutamine. These results might also indicate a decreased cellular influx of asparagine, since it is abundant in CM. Flux studies will be necessary to determine the exact relationship between glutamine synthesis and transport in PC cells upon receptor activation.

To explain the role of OR51E2 in PC, we propose the following model: agonist stimulation generates new cells through asymmetric division and gradually increases the subpopulation of terminally differentiated cells expressing neuroendocrine markers (see graphical abstract).

NE-like cells from PC are characterized by increased expression of NSE and AMACR and decreased expression of K18 and AR ^34,93,94^. Increased expression of NSE and AMARC and decreased expression of AR and K18 following 19-OH AD and AFMK treatment demonstrate that these OR51E2 agonists induce a neuroendocrine phenotype. We confirmed that this effect is receptor-mediated, as treatment of OR51E2-knockout LNCaP cells significantly reduced the NSE and AMACR transcript levels.

Cell proliferation and differentiation have an inverse relationship, and terminal differentiation coincides with proliferation arrest and exit from the division cycle ^95^. Our results demonstrate that agonist treatment during the first 3 days induces cell proliferation at a rate similar to control cells, but after 4 days the viability of these cells, as measured by ATP content, was significantly reduced. Our results also demonstrate that receptor activation results in a new subpopulation of cells that undergoes G0/G1 cell cycle arrest and has decreased DNA synthesis, which is concordant with the results from our metabolomics analysis. Cellular senescence is an irreversible growth arrest, and senescent cells actively suppress apoptosis ^96^. We found that agonist treatment decreases the fraction of apoptotic cells, indicating that growth arrest likely induces cellular senescence. Future studies are needed to confirm the irreversibility of this process.

Furthermore, recent whole-exome sequencing of NEPC and CRPC showed an overlap in genomic alterations, and in both demonstrated increased amplification of the OR51E2 gene, supporting our hypothesis that this receptor contributes to the NE-phenotype of PC ^79,80^.

Prostatic adenocarcinomas typically contain foci of non-proliferating NE-like cells that increase in number as cancer progresses ^97^. Although these cells are non-mitotic, proliferating carcinoma cells have been found in their proximity, suggesting that the non-proliferating NE-like cells likely provide paracrine stimuli for growth of the surrounding carcinoma cells ^98,99^. Our results demonstrate that activation of OR51E2 by newly-discovered PC-relevant agonists induces and/or facilitates cellular transformation, resulting in NEtD, a characteristic phenotype of CRCP. This indicates that activation of OR51E2 in PC might contribute to development of non-proliferating foci. Our data demonstrate that activation of OR51E2 results in NEtD and establish this GPCR as a novel and therapeutic target for NEPC and CRPC.

## Author Contributions

Conceptualization, T.A.; Methodology, T.A., J.B., M.J.M., I.S. and S.Y.K.; Investigation, T.A., J.B., M.J.M., S.Y.K., I. S. and S. L.; Writing, T.A.; Writing, Review, and Editing, T.A., J.B., M.J.M., S.L., and H.M.; Resources, H.M.; Supervision, T.A. and H.M.

## Acknowledgements

We would also like to thank Professors J. Heitman, J. Huang, M. P. Dewhirst and J. A. Chi, and Drs. C. de March and T. Zhou for their critical review on the manuscript. This study was supported by the NIDCD to H. Matsunami. The authors declare no conflict of interest.

## MATERIAL AND METHODS

### Homology Modeling

Amino acid sequences of bovine rhodopsin, the human adrenergic beta-2-receptor β 2AR, the mouse olfactory receptor MOR42-3, and the human olfactory receptor OR51E2, were initially aligned using a MAFFT program ^100^. Our model was built based on the crystal structure of β2AR (4LDO; ^101^). We used BioEdit Sequence Alignment Editor (http://www.mbio.ncsu.edu/BioEdit/bioedit.html) to manually remove gaps in the β2AR and OR51E2 sequences and re-align Cys178 in OR51E2 (EC2) with Cys191 from the β2AR sequence (**Figure S1**). β2AR has a disulfide bond between the conserved Cys106 residue at the N-terminal end of TM3 and Cys191 from EC2. Thus, to introduce a disulfide bond we re-aligned equivalent cysteines in OR51E2 (Cys96-Cys178). In addition, the most conserved residues were also aligned (Asp41, Leu54, Cys96, Leu114, Asp120, Arg121, Tyr122, Pro128, Pro159, Tyr217, Asp287, Pro288, Ile290, and Tyr290). We used Modeller v.9.14 to create homology models ^102^ (http://salilab.org/modeller/), and 20 homology models were made using an automodel script with the default optimization and refinement. Each model was assessed using a DOPE score^103^. The best model, which had a DOPE score of -37275, was chosen for further analysis. This .pdb file was imported into ICM Software (MolSoft v.3.8, LLC, La Jolla, CA) ^104^, and the protein structure was analyzed using Ramchandran Plot Analysis. Residues that were out of place (Glu90, Arg222, His180, Leu81, and Lys294) were optimized. The model was transformed into an ICM object and subjected to regularization and minimization using MMFF Cartesian minimization (300 steps). The minimized model was imported into Modeller and again evaluated to calculate DOPE score (new DOPE = −38897). This final, minimized model was used for docking and VLS in the ICM program.

### Virtual Library Screening

The library used for VLS was selected from the HMDB (www.HMDB.ca) and consisted of metabolites detected in human blood, tissue, urine, and saliva. VLS was performed with the ICM software as described previously ^69^. Briefly, the potential energy maps of the receptor were made in a box with a 0.5-Å grid and a size of 53 × 50 × 45 Å. The initial position of the ligand was set within the center of this box, which extends to the receptor interior (above the center positions of Ser107 and Ala108, which are equivalent positions to Ile112 and Val113 in MOR42-3 that are known to be part of the ligand binding pocket ^105^). The docking stimulation was set to 1 and the maximal number of conformations to 10. The remainder of the docking parameters were set at the default values. The VLS result lists ligand-metabolite pairs according to their scores, and lower scores indicate ligands that are more likely to bind to the receptor. Binding of the ligand to the receptor is described using Score and mfScore functions. The Score function is based on a *theoretical calculation* of receptor-ligand binding energy:

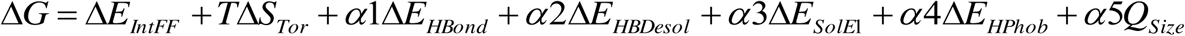

The mfScore function is a *knowledge-based* potential derived from the frequencies of occurrence of various atom pairs within the experimental ligand/receptor complex structures deposited in the Protein Data Bank (PDB) ^106^. It represents a measure of statistical probability of interaction between the ligand and receptor. Our previous results indicate that both scoring functions were equally successful in predisting ligands ^69^. The top 50 hits from each scoring list are presented in **Tables S1 and S2**.

### Cell Culture

Prostate cancer LNCaP-FGC cells derived from lymph node metastatic site (passage 30–32) were obtained from the American Type Culture Collection (ATCC). Cells were maintained at 37°C in RPMI-1640 medium (Sigma R8758) enriched with 0.5% glucose (45%, Sigma G8769), 1% 1 M HEPES (Gibco-Thermo 15630), 1% 100 mM Na-pyruvate (Gibco-Thermo 11360), 10% FBS (HyClone, SH30071.03), 100 U/mL penicillin, and 0.1 mg/mL streptomycin. At 4 to 5 days after seeding, cells were 70% to 80% confluent and the stimulus was applied at the indicated concentration for the indicated time. The medium was replaced every fourth day.

### Luciferase Assay

Briefly, the OR51E2 plasmid was transfected into HANA3A cells along with a CREB-dependent luciferase (firefly) and a constitutively active luciferase (*Renilla*) ^46^. Upon ligand binding, an increase in cAMP drives the expression of firefly luciferase and increases the signal. To control for variation in cell number and transfection efficiency, the luciferase signal was normalized to the activity of the *Renilla* luciferase in the same cells. All stock solutions of chemicals were prepared either in dimethylsulfoxide (DMSO) or ethanol. Final dilutions were made in M10d medium. M10d is MEM medium enriched with 5% dialyzed FBS serum, which is devoid of small molecular weight compounds (<10,000 Da), since OR51E2-transfected cells gave a high luciferase signal in the CD293 medium (Gibco 11913-019, supplemented with 30 μM CuCl_2_ and 2 μM L-glutamine) even in the absence of chemical stimulation and when compared to the basal activity of the control OR2W1 receptor-expressing cells (data not shown). All compounds that did not show agonist activity in M10d were later diluted in CD293 medium and tested for antagonist activity. The rest of the protocol was performed as previously published ^46^. Cells were exposed to candidate ligands for 3.5 hours at various concentrations. For each compound that showed a response >2 SD of the baseline response (no chemical applied), the EC50 or IC50 was determined from a sigmoid dose-response curve using a Graph-Pad Prism (Graph-Pad Prism Software, San Diego, USA). Data were fitted to the equation: Y = Bottom + (Top-Bottom)/(1+10^^^((LogIC50-X) *HillSlope)).

### 19-OH AD measurements using LC/MS

LNCaP cells were first grown in T-75 flasks until just fully confluent. Cells were then split into 6 T-25 flasks Cells were exposed to 250 μM AFMK, or to the medium only, for 3 days. We used phenol-red free RPMI 1640 with 10% CD-serum (Hyclone, SH30068) or RPMI 1640 with 10% FBS, as described in cell culture protocol. After 3 days, medium was collected and frozen at −80^o^C until LC/MS measurements.

In 2 mL polypropylene vial, 800 μL of cell media sample was vigorously mixed with 1 mL of ethyl acetate, centrifuged, and 900 μL of the organic (top) layer evaporated under a stream of nitrogen at room temperature. To the dried residue, 20 μL of the following reagent for picolinoyl ester derivatization was added: 40 mg of picolinic acid, 20 mg of 2-methyl-6-nitrobenzoic anhydride (MNBA), 10 mg of 4-dimethyl-laminopyridine, 1 mL acetonitrile, and 20 μL trimethylamine. After 20 min incubation at 50^o^C, 30 μL of 0.1% formic acid in water was added and 25 μL sample injected into the LC-MS/MS system (slightly modified from ^107^. LC conditions (Shimadzu 20A series HPLC): Agilent Eclipse PLUS, C18, 50 x 4.6 mm, 1.8 μm particle size column; mobile phases A/B: 0.1% formic acid in water/acetonitrile; elution gradient: 0–1min 20–90%B, 1–1.5min 90%B, 1.5–1.7min 90–20%B; runtime: 5min. Mass spectrometer conditions (AB/Sciex API5500 QTrap): after optimization of the electrospray and quadrupoles parameters by infusion of 100 ng/mL 19-OHAD at 10 μL/min rate, 408.2/267.2 MS/MS transition was used for quantification.

### Metabolomics GC-MS Analysis

LNCaP cells were first grown in T-75 flasks until just fully confluent. Cells were then split into 6 T-25 flasks and exposed to agonists the following day: 250 μM AFMK, 100 nM 19-OH AD, or 1 mM propionic acid (PA), with 6 biological replicates in each group. After agonist treatment for 3 days, CM was removed and banked at −80°C, and the cells were rinsed with 10 mL ice-cold PBS. Next, 3 mL of ice-cold 0.9% NaCl were added, and cells were scraped off of the plates and transferred to 5-mL tubes that were previously cleaned with acetonitrile. Each flask was rinsed with an additional 2 mL of ice-cold 0.9% NaCl, and the cell suspensions were transferred to the 5-mL tubes and centrifuged at 1000 rpm for 5 min at 4°C. Supernatant was removed, and 200 μL of ice-cold dH_2_0 with 0.6% formic acid were added to the cell pellets. Then, 20 μL of resuspended cell pellets were separated for protein measurements (Qubit Protein Assay kit, Cat. # Q33211, Thermo Fisher Scientific), and 180 μL of acetonitrile were added to the remaining pellet. Prepared cell lysates were maintained at −80°C until metabolomics analysis.

For exploratory, non-targeted metabolomics via gas chromatography/mass spectrometry (GC/MS), metabolites were extracted from cell lysates with methanol, methoximated in dry pyridine, and then silylated with *N*-methyl-*N*-(trimethylsilyl) trifluoroacetamide. Samples were analyzed on a 6890N GC connected to a 5975 MS (Agilent Technologies, Santa Clara, CA) equipped with 2 wall-coated open-tubular (WCOT) GC columns connected in series (Agilent part 122–5512, DB5MS, 15 m in length, 0.25 mm in diameter, with an 0.25-μm luminal film) separated by a microfluidic flow splitter to enable hot back-flushing at the end of each run ^108^. Data were acquired by scanning from *m/z* 600 to 50 as the oven ramped from 70°C to 325°C. Data were deconvoluted using AMDIS software ^109^. Metabolites were identified using our retention time-referenced spectral library ^110–112^, which is based in part on that of Kind *et al.* ^113^. Reported data are log_2_ transforms of the areas of deconvoluted peaks.

Data were normalized to the protein content in each sample. MetaboAnalyst 3.0 was used for statistical analysis ^114^. Briefly, peak intensity data were presented in columns and log_2_-normalized. We used unpaired analysis, and data were auto-scaled (mean-centered and divided by the standard deviation of each variable). For pathway analysis, we used a Globaltest pathway-enrichment analysis algorithm ^115^ in MetaboAnalyst 3.0.

### RT-PCR Analysis

LNCaP cells were grown in T-75 flasks until fully confluent. A split ratio of 1:6 was used to subculture the cells in T-25 flasks for 24 to 48 hours before stimulation. Stimulation with agonists lasted 3 or 12 days. Medium was changed every 4 days in the 12-day experiment. Total RNA was extracted using Trizol reagent (Invitrogen) and cleaned using the RNeasy kit (Qiagen) following the manufacturer’s protocol. Integrity of total RNA was confirmed by agarose gel electrophoresis. cDNA was generated by reverse transcription using SuperScript II Reverse Transcriptase (Invitrogen). Primers for the following genes were designed with the Primer 3.0 program: OR51E2, NSE-neuron-specific enolase, AMACR-α-methylacyl-CoA racemase, a1H T-type calcium channel (Cav3.2), AR-androgen receptor, GAPDH, keratin 5, keratin 8, and keratin 18 (Table S3). PCR amplification was performed with HotStart Taq Polymerase (Qiagen) using the following protocol: 95°C for 15 min followed by 30 cycles at the following times and temperatures: 95°C for 15 s, 55°C for 15 s, and 72°C for 30 s. Final annealing was done at 72°C for 5 min. Expression levels of the test genes were normalized to GAPDH.

### Cell Proliferation Assay

Cells were plated in five 96-well plates (100 μL cell suspension per well) and after overnight attachment when cells were 15% to 20% confluent, cells were stimulated with increasing concentrations of selected agonists. Cell growth and viability was estimated each day for the next 4 days by using Cell Titer-Glo Luminescent Cell Viability Assay (Promega, Cat. No. G7570) according to the manufacturer’s instructions.

### Cell Cycle Analysis

LNCaP cells were plated in 100-mm Petri dishes. After overnight attachment, when cells were 40% to 50% confluent, cells were stimulated once with 100 nM 19-OH AD and 250 μM AFMK for the next 4 days. Cells incubated in the regular RPMI-1640 medium served as controls. After treatment, cells were trypsinized with 0.05% trypsin-EDTA (Gibco) and fixed with 70% ethanol in PBS for 15 min at 4°C. Fixed cells were washed and incubated with 50 μg/μL RNase A and 20 μg/μL propidium iodide and subjected to cell cycle analysis using a Becton Dickinson FACSCalibur cytometer. Data were analyzed with BD CellQuest software. Experiments were performed 3 times in triplicate (3 biological and 3 technical replicates).

### LNCaP OR51E2 Knockout Cell Lines

To generate CRISPR/Cas9-mediated knockout LNCaP cells, we cloned sgRNAs targeting exon 2 of OR51E2 into LentiCRISPR.v2 (Addgene #52961) for coexpression of sgRNA with *Streptococcus pyogenes* Cas9. We prepared lentiviral particles for each sgRNA vector by cotransfecting HEK293T cells with the LentiCRISPR vectors psPAX2 and pMD2.g using TransIT-LT1 (Mirus). LNCaP cells were transduced with this virus at a multiplicity of infection (MOI) of <1 in the presence of 8 μg/ml polybrene. At 24 hours post-infection, cells were selected with 2 μg/mL puromycin for 72 hours and then expanded for verification of gene editing and experimental analysis. Cells were harvested for genomic DNA and PCR amplification of the OR51E2 locus for the Surveyor Assay (Integrated DNA Technologies) 1 week after infection. Additionally, for stable cells harboring OR51E2 sgRNA #1, PCR products were cloned into a TOPO-TA cloning vector (ThermoFisher Scientific) and Sanger sequenced to assess the rate of mutation.

Sequences:

sgRNA #1:CGTGGTCTTCATCGTAAGGA

sgRNA #2:AGGCCTCAAAGCTAATCTCT

sgRNA #3:CATT GAAT CCACCATCCTGC

OR51E2 forward:ACGAAGGTATGGACCAGTAGGA

OR51E2 reverse:AAGACCATATACCACATTGGGC

### Quantification and Statistical Analysis

All results are expressed as means ± SEM. Computations assumed that all groups were samples from populations with the same scatter. The investigators involved in this study were not completely blinded during sample collection or data analysis. Significance was determined by (multiple) two-tailed unpaired t-test using Prism 7 software. A P value of 0.05 was considered significant.

### Reagent and Resource Table

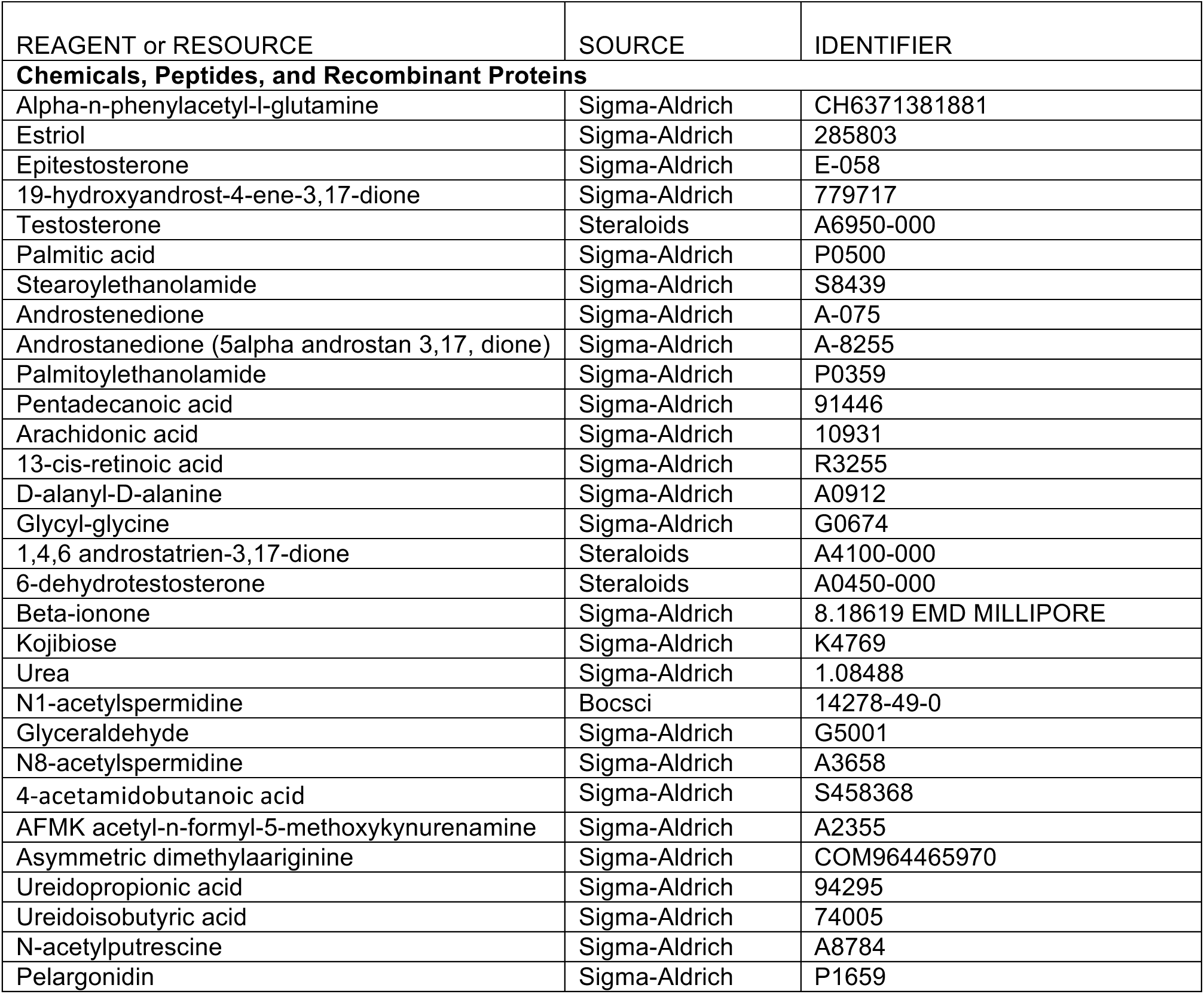
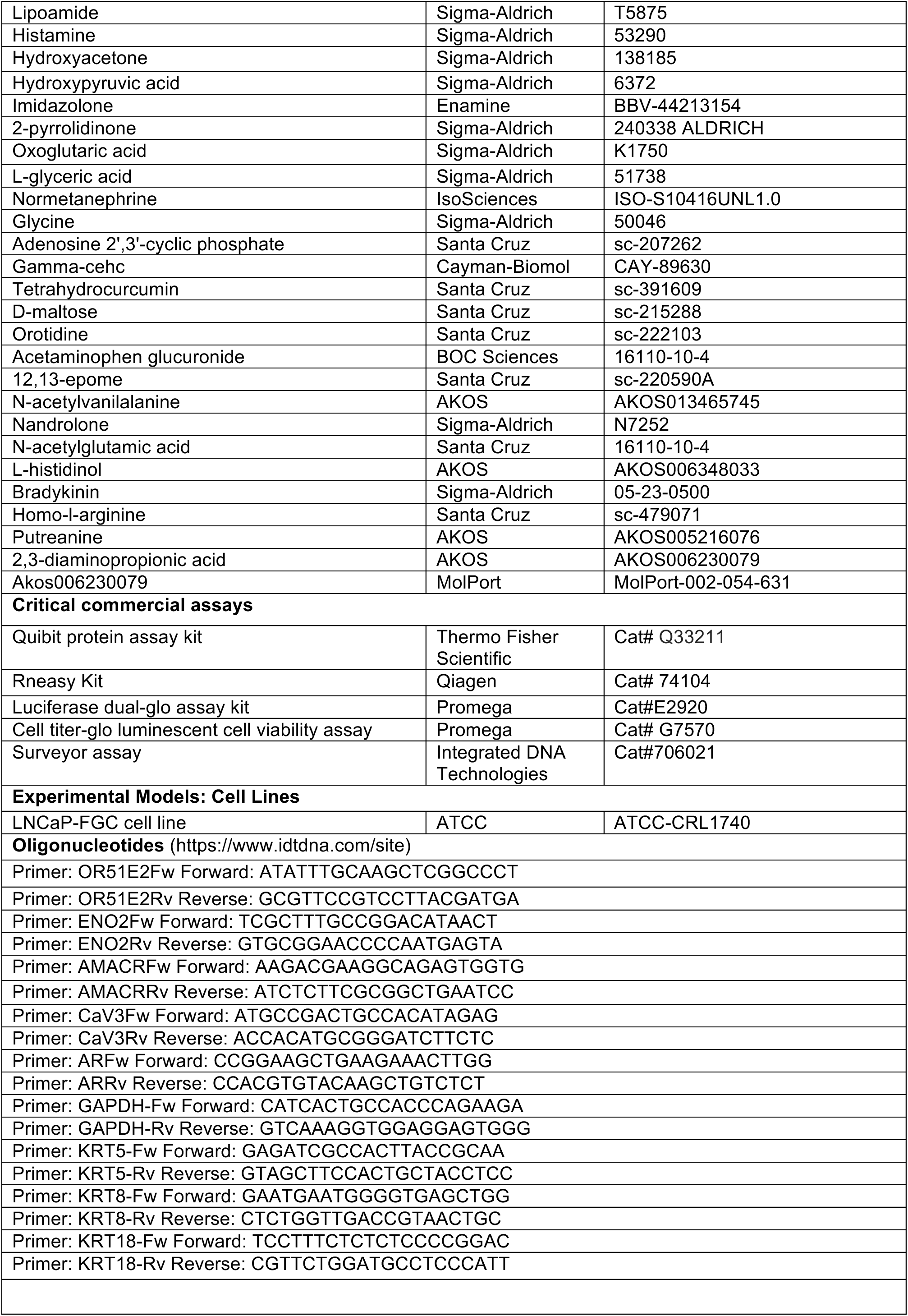
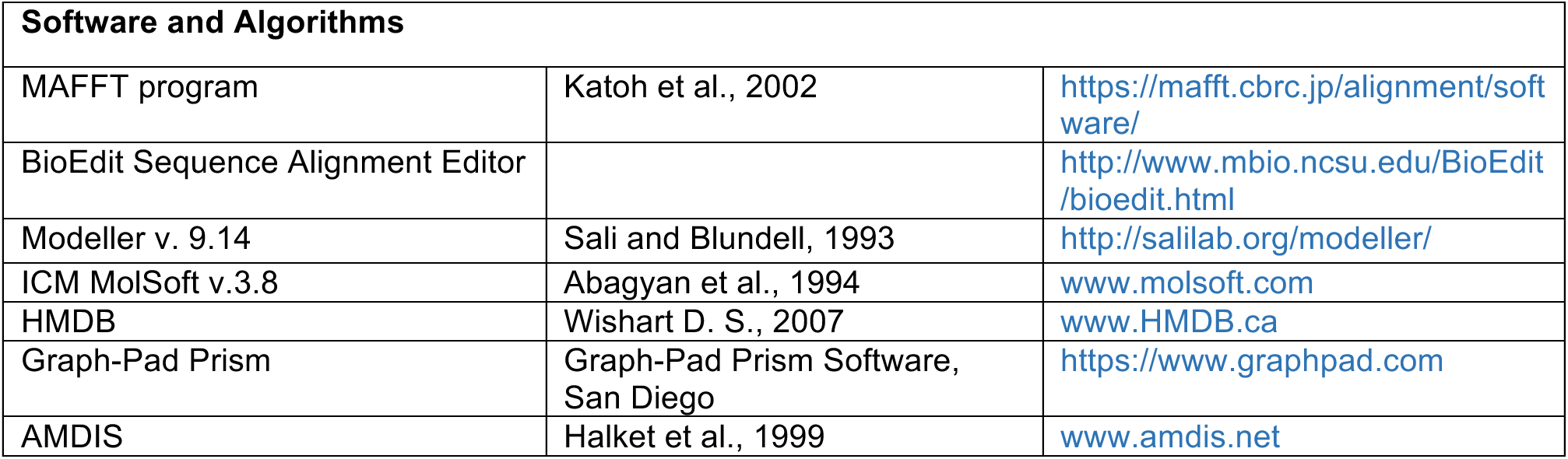

